# Caveolin-1 mediates neuroinflammation and cognitive impairment in SARS-CoV-2 infection

**DOI:** 10.1101/2023.10.18.563024

**Authors:** Troy N. Trevino, Avital B. Fogel, Richard Minshall, Justin M. Richner, Sarah E. Lutz

## Abstract

Leukocyte infiltration of the CNS can contribute to neuroinflammation and cognitive impairment. Brain endothelial cells regulate adhesion, activation, and diapedesis of T cells across the blood-brain barrier (BBB) in inflammatory diseases. The integral membrane protein Caveolin-1 (Cav-1) critically regulates BBB permeability, but its influence on T cell CNS infiltration in respiratory viral infections is unknown. In this study, we sought to determine the role of Cav-1 at the BBB in neuroinflammation in a COVID-19 mouse model. We used mice genetically deficient in Cav-1 to test the role of this protein in T cell infiltration and cognitive impairment. We found that SARS-CoV-2 infection upregulated brain endothelial Cav-1. Moreover, SARS-CoV-2 infection increased brain endothelial cell vascular cell adhesion molecule-1 (VCAM-1) and CD3+ T cell infiltration of the hippocampus, a region important for short term learning and memory. Concordantly, we observed learning and memory deficits. Importantly, genetic deficiency in Cav-1 attenuated brain endothelial VCAM-1 expression and T cell infiltration in the hippocampus of mice with SARS-CoV-2 infection. Moreover, Cav-1 KO mice were protected from the learning and memory deficits caused by SARS-CoV-2 infection. These results indicate the importance of BBB permeability in COVID-19 neuroinflammation and suggest potential therapeutic value of targeting Cav-1 to improve disease outcomes.

## Introduction

COVID-19 is associated with leukocyte infiltration of the CNS (Fernández-Castañeda et al., 2022; Lee et al., 2021; Lee et al., 2022; Schwabenland et al., 2021; Soung et al., 2022; Thakur et al., 2021). Some data suggests that T cells in the CNS contribute to neuroinflammatory processes in COVID-19 (Heming et al., 2021; Lee *et al*., 2022); for example, high perivascular T cell density spatially correlates with microglial nodules enriched in disease-associated proinflammatory phenotypes (Schwabenland *et al*., 2021). Immunopathologic contribution of T cell infiltration to microglial activation and neurodegeneration is commonly observed in CNS diseases (Ai and Klein, 2020; Cruz-Orengo et al., 2014; Dong and Yong, 2019). Hippocampal T cell infiltration particularly contributes to cognitive impairment, consistent with the important role of the hippocampus in cognition (Garber et al., 2019). Disorders of cognition are frequent in COVID-19 (Spudich and Nath, 2022). The contribution of hippocampal T cell infiltration to cognitive impairment in COVID-19 is incompletely understood.

Blood brain barrier (BBB) endothelial cells regulate T cell infiltration of the CNS. One way to address how T cell interactions at the BBB influence cognitive outcomes in COVID-19 is to assess function of specific proteins. Here, we focused on Caveolin-1 (Cav-1) (Jones and Minshall, 2020; Ohi and Kenworthy, 2022; Parton, 2018). Cav-1 promotes BBB permeability by facilitating endocytosis and transcellular transcytosis (Andreone et al., 2017; Ayloo and Gu, 2019; Chow and Gu, 2017; Liebner et al., 2018; Pandit et al., 2020; Pol et al., 2020). Adhesion molecules are essential for the adhesion between leukocytes and endothelial cells that ultimately leads to diapedesis into inflamed tissues (Martin-Blondel et al., 2015; Soldati et al., 2023). Cav-1 contributes to the scaffolding, membrane retention, and recycling of leukocyte adhesion molecules including vascular cell adhesion molecule (VCAM)-1 and Intracellular Adhesion Molecule (ICAM)-1 (Carman and Springer, 2004; Dragoni et al., 2017; Dragoni et al., 2020; Eum et al., 2015; Han et al., 2010; Lolo et al., 2022; Millan et al., 2006). Cav-1 upregulation in disease contributes to BBB leakage and neuroinflammation (Errede et al., 2012; Knowland et al., 2014; Lutz et al., 2017; Wu et al., 2016; Yang et al., 2020; Zhang et al., 2022). Indeed, we and others have shown that Cav1 promotes migration of proinflammatory T cells across the BBB, whereas loss of endothelial Cav-1 impairs T cells and neutrophil adhesion and transmigration (Hu et al., 2008; Lutz *et al*., 2017). Moreover, suppressing Cav-1 reduces proinflammatory leukocyte infiltration, neuroinflammation, and neurodegeneration (Lutz *et al*., 2017; Wu *et al*., 2016; Xu et al., 2013; Zhang *et al*., 2022).

Cav-1 might contribute to neuroinflammation in COVID-19. Strikingly, Cav-1 is upregulated in forebrains of COVID-19 decedents (Green et al., 2022; Premkumar and Sajitha Lulu, 2023). Transcellular BBB permeability is described in models of COVID-19 (Krasemann et al., 2022; Rhea et al., 2021; Zhang et al., 2021). These data raise the possibility that cerebrovascular Cav-1 upregulation might contribute to COVID-19 neuropathogenesis. However, the extent to which Cav-1 contributes to neuroinflammation and cognitive impairment in COVID-19 has not been tested.

Thus, the goal of the present study was to evaluate the extent to which Cav-1 contributes to cognitive impairment by promoting BBB permeability to T cells in a COVID-19 mouse model. We found that mild respiratory SARS-CoV-2 infection increased expression of Cav-1 and VCAM-1 on brain endothelial cells. This was accompanied by T cell neuroinflammation in the hippocampus and learning/memory deficits in infected mice. Genetic deficiency in Cav-1 offered protection from SARS-CoV-2 induced neuroinflammation and memory deficits. These data indicate that Cav-1-mediated BBB permeability to T cells is increased during acute SARS-CoV-2 respiratory infection and may contribute to neuropathology and cognitive impairment in COVID-19.

## Results

To test if Cav-1 is altered at the BBB during COVID-19, we infected 12-month-old C57Bl/6 (WT) mice with SARS-CoV-2 strain MA10 (Dinnon et al., 2020; Leist et al., 2020). This led to viral replication in the lung and mild respiratory disease (Dinnon *et al*., 2020; Leist *et al*., 2020). 12-month-old mice were used to recapitulate the more pronounced neuroinflammation observed in middle- and advanced-age COVID-19 patients (Amruta et al., 2022; Davis et al., 2022). At 5 days-post-infection (DPI) mice were euthanized and Cav-1 expression in brain microvessels was assessed by immunostaining (**Figure 1A-C**). We found that SARS-CoV-2 infection significantly increased Cav-1 positive area in the hippocampus, with a markedly perivascular distribution (**Figure 1C**). This suggested that brain endothelial Cav-1 is upregulated by SARS-CoV-2. We confirmed these results by quantifying Cav-1 by flow cytometry on brain endothelial cells acutely isolated from mice at 5DPI. The percentage of brain endothelial cells with high Cav-1 expression was increased by SARS-CoV-2 infection (**Figure 1D-G**). These data indicate that infection with SARS-CoV-2 upregulates Cav-1 expression in brain endothelial cells.

**Figure 1.**
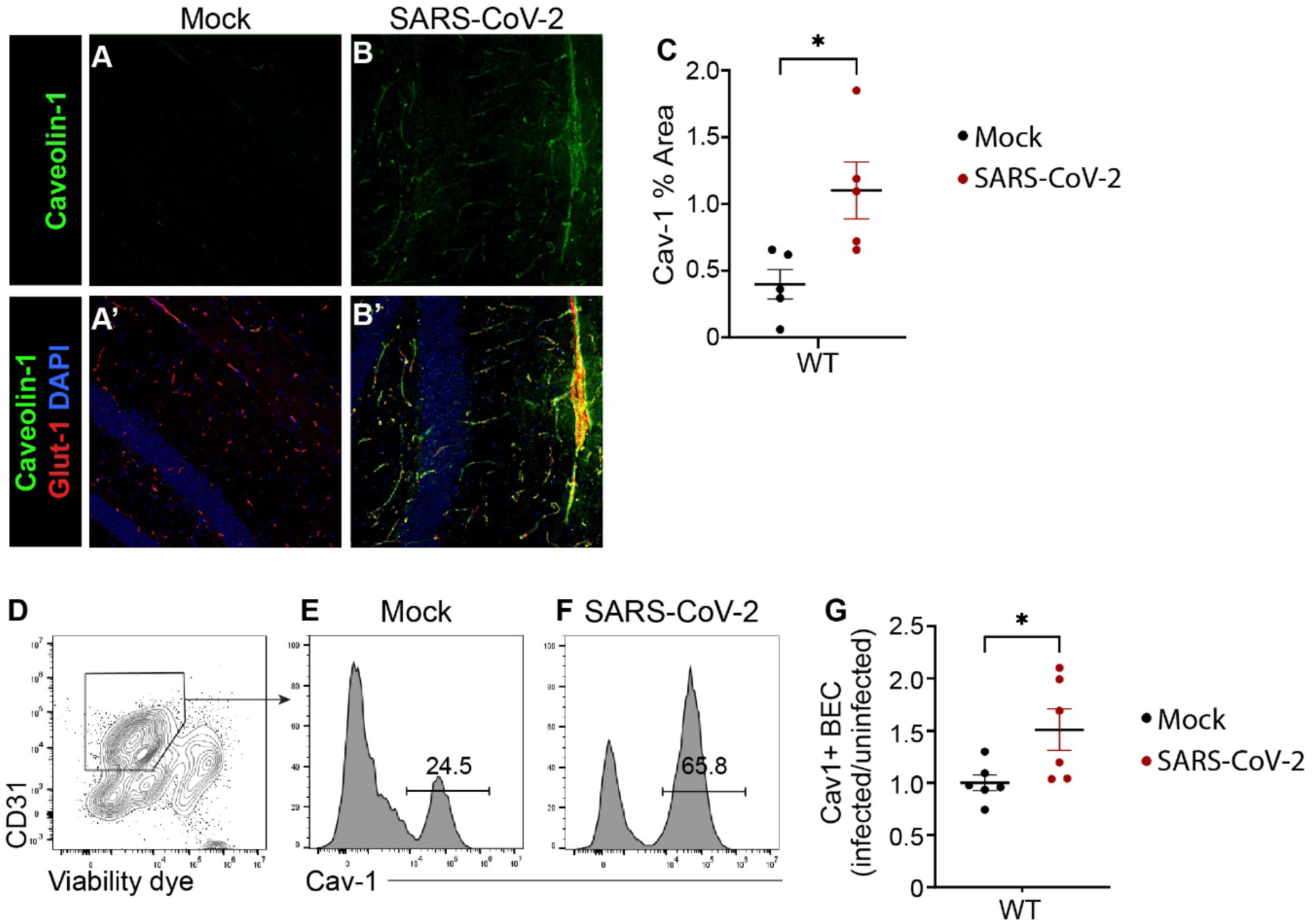
Cav-1 is increased in brain endothelial cells in SARS-CoV-2 infection. A-B) Immunofluorescence detection of Cav-1 (green) in hippocampal sections from wild-type mice euthanized 4 days after respiratory inoculation with SARS-CoV-2. Monochromatic images (A, B) and overlays between Caveolin-1 (green), brain endothelial cell protein Glut-1 (red), and DAPI (blue) (A’, B’). C) Quantification of % area immunoreactive for Cav-1 in hippocampus sections. n=3 mice/group, with 3 sections analyzed per mouse. p<0.05, unpaired student t-test. D) Flow cytometric plot of CD31 and viable exclusion dye demonstrates gating strategy for endothelial cells (box) in single cell suspension prepared from isolated brain microvessels of WT mice euthanized 4 days after respiratory inoculation with SARS-CoV-2. E-F) Histograms of Cav-1 fluorescence intensity in viable CD31+ endothelial cells isolated as in D. G) Quantification of Cav-1 fluorescence intensity in viable CD31+ endothelial cells from mice euthanized at 4 days after respiratory inoculation with SARS-CoV-2, expressed as ratio to healthy WT. n=6 mice/group from two independent experiments. p<0.01, unpaired student t-test.

We next analyzed VCAM-1 expression on brain endothelial cells isolated from SARS-CoV-2 infected wild-type mice. The percentage of brain endothelial cells highly expressing VCAM-1 was significantly increased in MA10 SARS-CoV-2 infected mice (**Figure 2A-B**). Endothelial Cav-1 on inflamed endothelial cells colocalizes with ICAM-1 and VCAM-1 (Hu *et al*., 2008). Moreover, loss of Cav-1 impairs adhesion and transmigration of T cells and neutrophils (Hu *et al*., 2008; Lutz *et al*., 2017; Wu *et al*., 2016). Therefore, we interrogated neuroinflammation during SARS-CoV-2 infection in mice with Cav-1 genetic deficiency (Cav-1 KO). VCAM-1 expression on brain endothelial cells was analyzed by flow cytometry. Brain endothelial cells from Cav-1 KO mice failed to upregulate VCAM-1 in response to SARS-CoV-2 infection (**Figure 2A-B**). This indicates that Cav-1 contributes to brain endothelial cell immune activation in SARS-CoV-2 infection.

**Figure 2.**
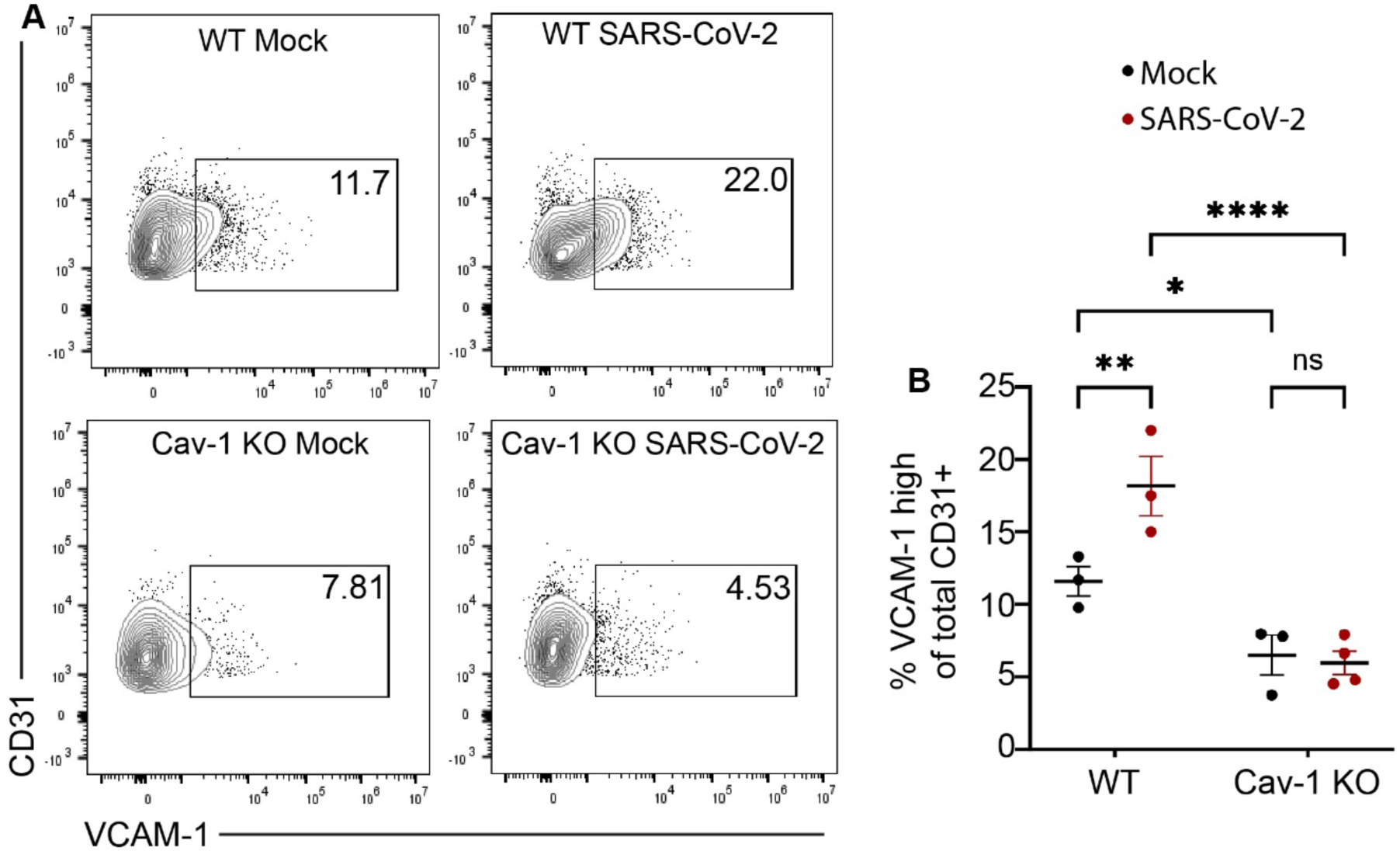
Cav-1 deficiency reduces brain endothelial cell VCAM-1 expression in SARS-CoV-2 infection. A) Flow cytometry of VCAM-1 and CD31 in brain endothelial cells. B) Quantification of VCAM-1^high^ cells as a percent of total CD31+ viable cells. n=3-4 mice/group. Two-way ANOVA demonstrated significant effect of genotype, disease, and genotype*disease interaction. Tukey’s multiple comparison tests revealed significantly increased BEC VCAM-1 in WT (p<0.01) but not Cav-1 KO mice with SARS-CoV-2. VCAM-1^high^ BEC were significantly fewer in Cav-1 KO mice with SARS-CoV-2 than in WT mice with SARS-CoV-2.

We next analyzed T cell infiltration into the hippocampal parenchyma in WT and Cav-1 KO mice infected with MA10 SARS-CoV-2. T cell neuroinflammation in the hippocampus was apparent in SARS-CoV-2 infected mice compared to healthy controls (**Figure 3**). Importantly, we observed fewer perivascular CD3+ T cells in the hippocampus in SARS-CoV-2 infected Cav-1 KO mice compared to infected WT mice (**Figure 3A-E**). This suggests that Cav-1 deficiency may offer protection against SARS-CoV-2 neuroinflammation by attenuating T cell adhesion to and migration across the BECs.

**Figure 3.**
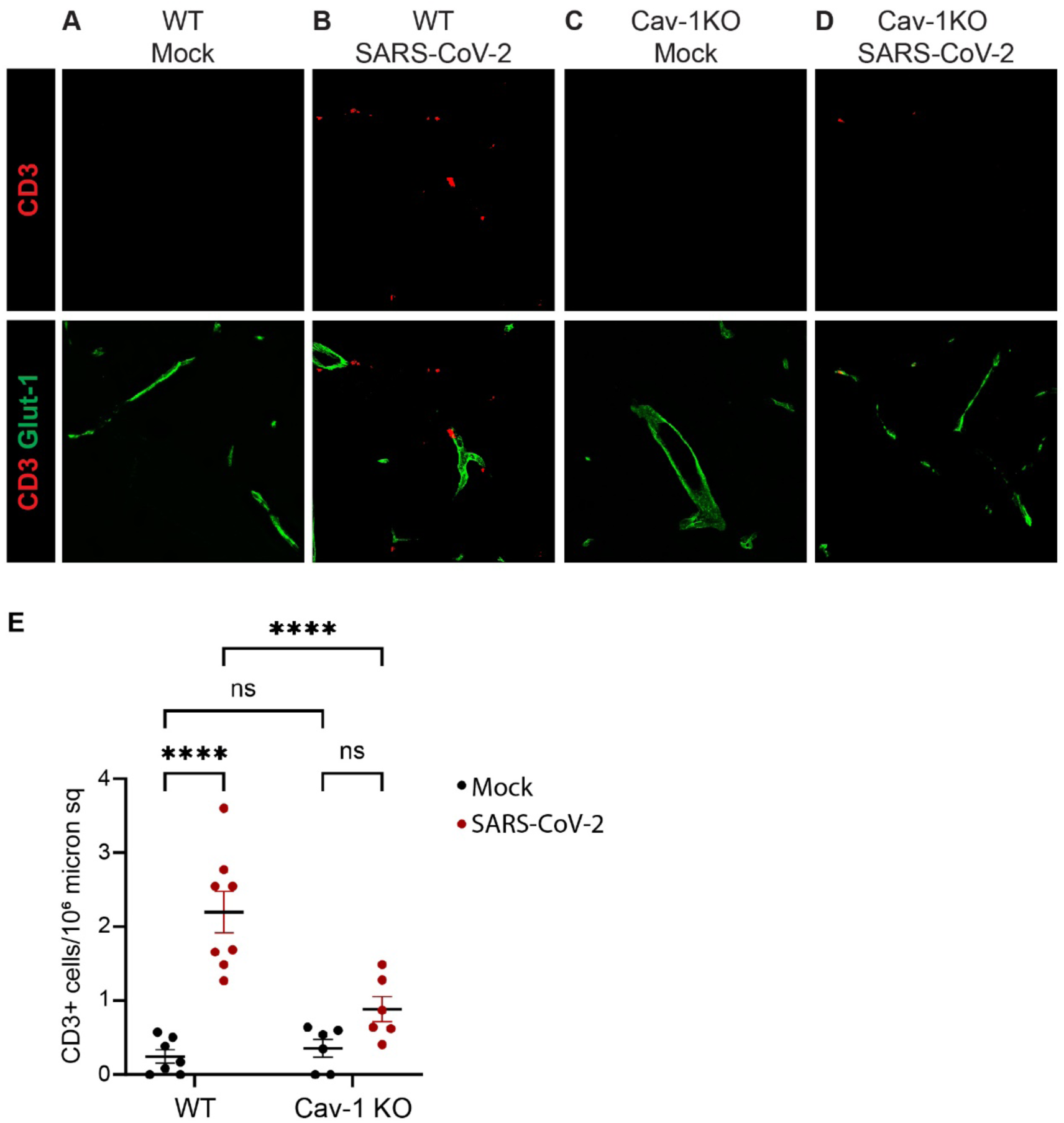
Cav-1 deficiency reduces hippocampal T cell density in SARS-CoV-2 infection. A-D) Immunofluorescence detection of CD3+ T cells (red) and Glut-1+ cerebrovasculature (green) in hippocampal sections of mice euthanized at 5 days after inoculation with SARS-CoV-2. E) Quantification of CD3+ T cells in hippocampal sections. n=6-8 mice/group, with two sections analyzed per mouse. Two-way ANOVA demonstrated significant effect of genotype, disease, and genotype*disease interaction. Tukey’s multiple comparison tests revealed significantly more T cells in hippocampus of WT mice with SARS-CoV-2 as compared with vehicle-inoculated WT mice (p<0.0001), and significantly fewer T cells in hippocampus of Cav1 KO mice with SARS-CoV-2 as compared to WT mice with SARS-CoV-2 (p<0.01).

Neurological impairment can accompany acute SARS-CoV-2 infection (Spudich and Nath, 2022). The extent to which BBB inflammation influences neurological deficits in COVD-19 is unclear. We therefore tested whether Cav-1 deficiency offers protection from SARS-CoV-2 induced cognitive impairment. We utilized a novel object recognition to measure learning and memory related to hippocampal function. Cav-1 KO mice have some age-dependent neurobehavioral abnormalities, including spatial learning deficit and center avoidance (Gioiosa et al., 2008; Trushina et al., 2006). Nonetheless, in our study, uninfected Cav-1 KO mice had similar novel object recognition as uninfected WT mice (**Figure 4A**). In WT mice, SARS-CoV-2 respiratory infection significantly impaired the ability to discriminate between known and unknown objects (**Figure 4A**). Importantly, Cav-1 deficiency mitigated this effect of SARS-CoV-2 on learning and memory (**Figure 4A**). As expected, no object preferences were observed during the training task (**Figure 4B**). Together, these data suggest that Cav-1 upregulation on BECs may drive neuroinflammation and neurological deficits in COVID-19.

**Figure 4.**
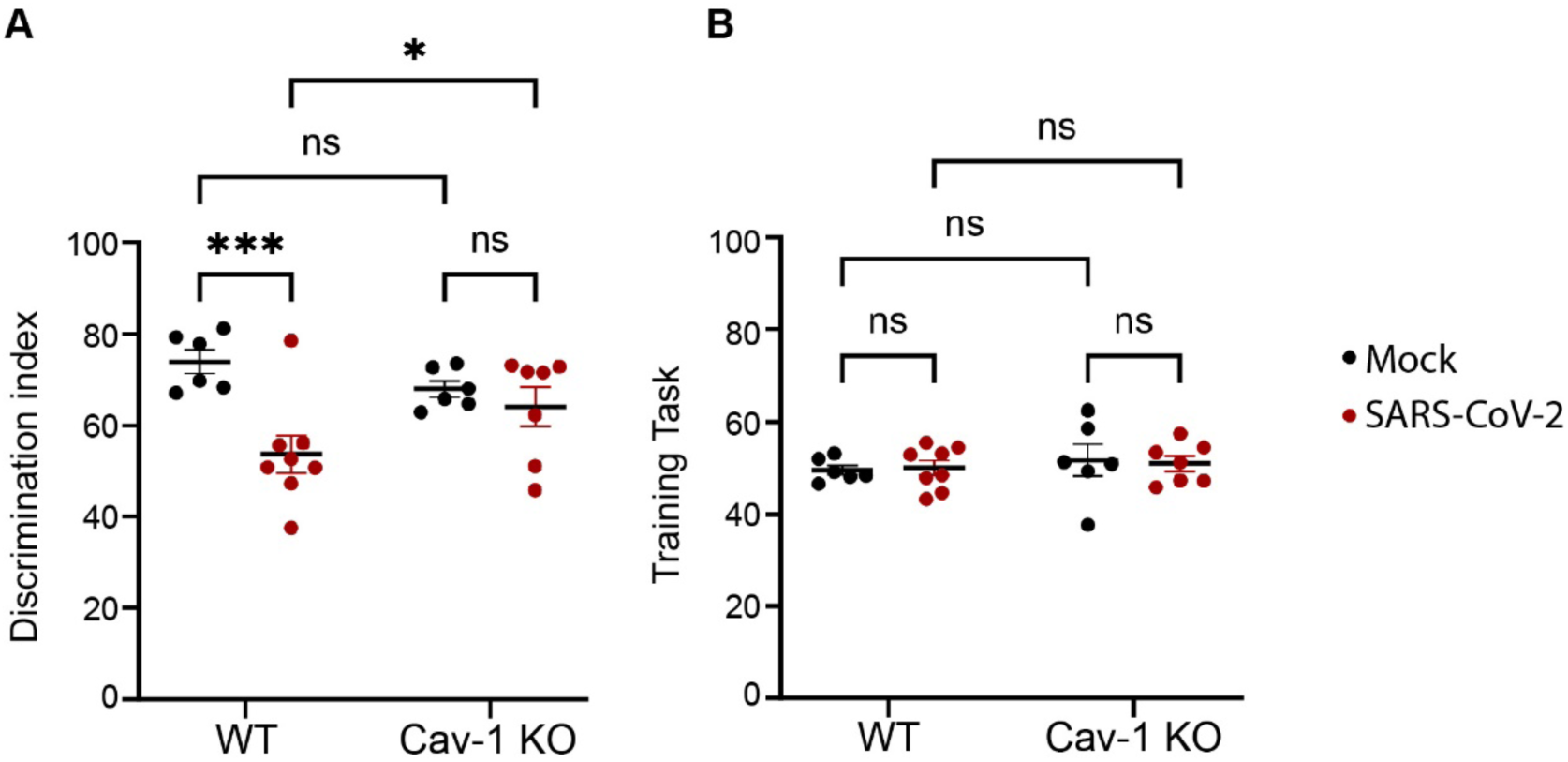
Cav-1 deficiency reduces short term learning and memory impairment in SARS-CoV-2 infection. A) Novel object discrimination index during novel object recognition testing at 4 days after inoculation with SARS-CoV-2. n=5-8 mice/group. Two-way ANOVA demonstrated significant effect of genotype, disease, and genotype*disease interaction. Tukey’s multiple comparison test revealed significant decrease in novel object discrimination in WT (p<0.001) but not Cav-1 KO mice with SARS-CoV-2. Cav-1 KO mice with SARS-CoV-2 performed significantly better than did WT mice with SARS-CoV-2. B) Object discrimination index measured during the training task demonstrates unbiased exploration during the training phase in all groups.

## Discussion

In this work, we sought to identify the role of brain EC dysfunction in SARS-CoV-2 induced neuroinflammation and cognitive impairment, and sought to determine whether deficiency in Cav-1 can offer protection. COVID-19-associated neurological impairment, cerebrovascular damage, and neuroinflammation are well-documented (Fernández-Castañeda *et al*., 2022; Hernández-Fernández et al., 2020; Hosp et al., 2021; Lee *et al*., 2021; Lee *et al*., 2022; Schwabenland *et al*., 2021; Soung *et al*., 2022; Spudich and Nath, 2022; Thakur *et al*., 2021). In this study, we observed that brain endothelial cell Cav-1 is upregulated by SARS-CoV-2 respiratory infection, indicating a potential role for Cav-1 and transcellular BBB permeability in COVID-19 neuroinflammation. This observation is consistent with a recent report of increased Cav-1 in the forebrain of COVID-19 decedents (Green *et al*., 2022). Our data suggest that enhanced brain endothelial Cav-1 may contribute to BBB leakage and neuroinflammation in COVID-19. Furthermore, we found that Cav-1 KO mice are partially protected from neurocognitive deficits associated with SARS-CoV-2 infection. These data suggest the potential therapeutic value of targeting Cav-1 mediated BBB permeability to improve disease outcomes.

Caveolin-1 can promote adhesion and migration of leukocytes across the BBB in part by influencing endothelial expression and distribution of leukocyte adhesion molecules (Knowland *et al*., 2014; Lutz *et al*., 2017; Song et al., 2007; Wu *et al*., 2016). We observed an increase in brain endothelial cell VCAM-1 in acute SARS-CoV-2 infection. This is similar to reports of upregulated leukocyte adhesion molecules in the cerebrovasculature of patients with severe acute COVID-19 (Savarraj et al., 2021; Thakur et al., 2023; Zhou et al., 2021) and in cultured brain endothelial cells exposed to SARS-CoV-2 virions or proteins (Buzhdygan et al., 2020; Constant et al., 2021; Motta et al., 2023; Yang et al., 2022). These findings fit within a larger context of brain endothelial hyperinflammatory responses and systemic inflammation in COVID-19 (Bonetto et al., 2022; Frere et al., 2022; Israelow et al., 2020; Pilotto et al., 2021; Rutkai et al., 2022; Soung *et al*., 2022). Here, we found that increased brain endothelial VCAM-1 correlates with increased Cav-1 expression in SARS-CoV-2 infection, and that VCAM-1 upregulation was attenuated in the absence of Cav-1. These observations indicate that Cav-1 contributes to cerebrovascular endothelial inflammation in COVID-19.

In neuroinflammation, brain endothelial cell VCAM-1 promotes leukocyte adhesion and migration into the CNS. Indeed, we found that enhanced Cav-1 and VCAM-1 on BECs from SARS-CoV-2 infected mice correlated with increased T cell infiltration into the hippocampus. Importantly, we observed T cell infiltration in the hippocampus was attenuated in Cav-1 KO mice with SARS-CoV-2. Perivascular leukocyte cuffing, microgliosis, and neuronophagia is observed in COVID-19 and its animal models(Amruta *et al*., 2022; Lee *et al*., 2021; Lee *et al*., 2022; Matschke et al., 2020; Schwabenland *et al*., 2021; Soung *et al*., 2022; Spudich and Nath, 2022; Thakur *et al*., 2021). The decreased hippocampal T cell density in the Cav-1 KO mice with SARS-CoV-2 is predicted to yield decreased microgliosis and neuronal damage. Interestingly, extensive perivascular leukocyte cuffing reportedly occurs only in a subset of COVID-19 decedents, suggesting that individual risk factors contribute to CNS presentation of infection (Agrawal et al., 2022; Thakur *et al*., 2021). Individual risk factors related to Cav-1 and other regulators of BBB adhesion and permeability may modify leukocytic infiltration and neurologic symptoms of SARS-CoV-2 infection (Adesse et al., 2022; Iadecola et al., 2020; Monje and Iwasaki, 2022; Teuwen et al., 2020; Vanderheiden and Klein, 2022; Vavougios et al., 2022).

Although our data supports central role for Cav1 in neuroinflammation and cognitive impairment, it is important to note caveats. In mice with Cav-1 deficiency in the entire body, protection from SARS-CoV-2 neuroinflammation might be due to pulmonary or cerebrovascular effects. By design, we focused on a phenotypic characterization of Cav-1. Due to the central role of Cav-1 in brain EC homeostasis, future studies are likely to identify numerous downstream processes by which loss of Cav-1 is protective in acute SARS-CoV-2 infection. Because SARS-CoV-2 is a BSL3 pathogen, we were limited in the kinds of neurobehavioral tests we could conduct to assess neuronal function, because of the practical limitations to conducting behavior tests within the confines of the biosafety cabinet in the BSL3 suite. This study exclusively examined acute infection, so we cannot draw conclusions regarding neuroPASC. Interestingly, however, neuroPASC correlates to neuroinflammatory cytokines (Fernández-Castañeda *et al*., 2022; Frere *et al*., 2022; Peluso et al., 2022) and serum markers of BBB leakage (Bonetto *et al*., 2022; Hanson et al., 2022). These observations suggest the value of future mechanistic studies into BBB Cav-1 dysregulation in neuroPASC.

## Methods

### Mice

All animal studies were approved by the UIC Animal Care and Use Committee (20-160, 21-051). Wild-type C57Bl/6 and Cav-1 KO (Jackson Laboratory 000664, 004585, respectively) mice were purchased from The Jackson Laboratory. Cav1 KO mice were backcrossed 9 generations by outbreeding to C57Bl/6J. For BioSafety Level 3 experiments, mice were housed on site in a specific pathogen free barrier suite for at least 7 days prior to initiation of experiments. Mice were transferred to the Animal BioSafety Level 3 facilities at least 2 days prior to inoculation. Mice were maintained on standard light-dark cycles with ad libitum food and water in micro-isolation cages.

### Behavioral Assays

Behavior tasks were conducted in a dim biosafety cabinet laminar flow hood in the BSL3 facility. Testing arenas were white plastic bins 13 inches x 19 inches (Ikea) with pebbled floor. For novel object recognition (NOR), we first tested a catalog of 10 objects for intrinsic preference. Objects were similar in size (1-2 inches wide, 3-4 inches tall), visually interesting, without smell, and made of easily cleaned non-porous materials, e.g. 25 ml suspension flasks filled with pebbles, 50 ml conical tubes filled with corncob bedding, or 3-D printed flagpoles. Neodynium magnets affixed to each object were used to ensure consistent object placement relative to magnets permanently affixed to the underside of the arena. For familiarization, we placed individual mice in an arena containing two suspension flasks and allowed 10 minutes exploration. Behavior was filmed with an overhead mounted wide-angle webcam (Logitech C920S HD Webcam). Intersession interval was 14 hours. For the testing session, mice were reintroduced into the arena containing one suspension flask and one novel object and filmed for 10 minutes. Objects and field were cleaned with ethanol and dried in between mice. Testing was conducted between 7:00-10:00AM. Videos were coded and independently scored by two blinded scientists for duration of exploration of each object. Preference was (sec investigating novel object)/(sec investigating any object)*100. 50% indicates no preference.

### Histology

Mice were euthanized 5 days after infection with SARS-CoV-2. Mice were perfused with PBS with a peristaltic pump. Brains were fixed in 4% paraformaldehyde and paraffin embedded. Antigen retrieval (10 mM Tri-sodium citrate [dihydrate], 0.05% Tween-20, pH 6.0; or Tris-EDTA pH 9.0 for Iba1) was 40 minutes at 98°C. Slides were blocked with 5% normal serum, 0.1% Triton-X 100. Primary antibodies incubated overnight at 4°C at 1:100 included CD3 (Cell Signaling 78588), Glut1 (Abcam ab40084), and Caveolin-1 (Invitrogen PA5-17447). Alexa-fluorophore conjugated secondary antibodies (Invitrogen) were incubated 2h at 22°C. Mounting medium contained DAPI (Ibidi 50011). Microscopy was conducted using Zeiss LSM880 or Leica DMI8 microscopes. Quantification was performed using FIJI (NIH).

### Endothelial cell isolation

Mice were transcardially perfused with PBS 5 days after infection. Brainstem microvascular endothelial cells were isolated using gradient centrifugation in 25% BSA (Marottoli et al., 2021). Microvessels were dissociated with collagenase/dispase (Millipore Sigma 10269638001) and DNase (Worthington LK003172) for 1 h in 37°C water bath and passed through 100 µm cell strainer (PluriSelect USA 43-10100-60). Dissociated microvascular cells were stained for flow cytometric analysis.

### Flow cytometry

Endothelial cells isolated from brain from SARS-CoV-2 infected mice were stained with fixable viability dye (Zombie Green, Biolegend 423111) and blocked with Fc receptor blockade (anti-mouse CD16/32, Biolegend 101319) followed by incubation with anti-CD31 antibody (BD Biosciences 561073) and anti-VCAM-1 antibody (Biolegend 105719). Fixation and permeabilization (Biolegend 421403) was followed with an antibody against Cav-1 (Cell Signaling 31411S). Stained cells were analyzed with a Beckman CytoFLEX S flow cytometer.

### Statistics

Statistics were conducted using GraphPad Prism 10. Pairwise comparisons used Student’s t-test, with Welch’s correction when variances were unequal. Grouped data were tested with two-way ANOVA; significant interactions were compared by Sidak’s multiple comparisons test.

## Acknowledgements

This work was supported by the DOD MS200290 and NIH KL2TR002002 (SEL), T32HL139439 (TNT), R01AI150672 (JMR), and University of Illinois Institutional Funds (SEL). Research support services were obtained from the Research Histology Core and the Center for Clinical and Translational Science Biostatistics Core at the University of Illinois at Chicago U54 TR002003.

